# Differential expression of core metabolic functions in *Candidatus* Altiarchaeum inhabiting distinct subsurface ecosystems

**DOI:** 10.1101/2023.11.20.567779

**Authors:** Sarah P. Esser, Victoria Turzynski, Julia Plewka, Carrie J. Moore, Indra Banas, André R. Soares, Janey Lee, Tanja Woyke, Alexander J. Probst

## Abstract

*Candidatus* Altiarchaea are widespread across aquatic subsurface ecosystems and possess a highly conserved core genome, yet adaptations of this core genome to different biotic and abiotic factors based on gene expression remain unknown. Here, we investigated the metatranscriptome of two *Ca*. Altiarchaeum populations that thrive in two substantially different subsurface ecosystems. In Crystal Geyser, a high-CO_2_ groundwater system in the USA, *Ca*. Altiarchaeum crystalense co-occurs with the symbiont *Ca*. Huberiarchaeum crystalense, while in the Muehlbacher sulfidic spring in Germany, an artesian spring high in sulfide concentration, *Ca*. A. hamiconexum is heavily infected with viruses. We here mapped metatranscriptome reads against their genomes to analyze the *in situ* expression profile of their core genomes. Out of 537 shared gene clusters, 331 were functionally annotated and 130 differed significantly in expression between the two sites. Main differences were related to genes involved in cell defense like CRISPR-Cas, virus defense, replication, and transcription as well as energy and carbon metabolism. Our results demonstrate that altiarchaeal populations in the subsurface are likely adapted to their environment while influenced by other biological entities that tamper with their core metabolism. We consequently posit that viruses and symbiotic interactions can be major energy sinks for organisms in the deep biosphere.

**(Originality-Significance Statement:** Organisms of the uncultivated phylum *Ca*. Altiarchaeota are globally widespread and fulfill essential roles in carbon cycling, *e*.*g*., carbon fixation in the continental subsurface. Here, we show that the transcriptional activity of organisms in the continental subsurface differ significantly depending on the geological and microbial setting of the ecosystem explaining many of the previously observed physiological traits of this organism group.)

## Main

The aquatic deep subsurface houses some of Earth’s most diverse and complex ecosystems, varying in chemical, physical and geological parameters. As a result, they serve as ecological niches for differently adapted microorganisms. DPANN (Diapherotrites, Parvarchaeota, Aenigmarchaeota, Nanohaloarchaeota and Nanoarchaeota) archaea^1^ are alongside with other archaea and bacteria found in various aquatic deep subsurface ecosystems such as marine and terrestrial geysers, lakes and boreholes^2–6^. To cope with their limited metabolic capacity, DPANN archaea are known to be able to live in symbiosis with other archaea, with the best studied example of *Ignicoccus hospitalis* and *Nanoarchaeota equitans*^2,7–9^, but also recently described DPANN-host associations from the branch of Micrarchaeota^10^. One exception to this rule is *Ca*. Altiarchaeum, a geographically widespread genus of organisms that live freely as carbon fixing organism in the deep subsurface. Recent investigations of *Ca*. Altiarchaea have shown site-specific genomic adaptations likely resulting from horizontal gene transfer, yet these organisms harbor a highly conserved core genome that follows a strict biogeographic pattern^11^.

This study focuses on metatranscriptomic expression profiles of *Ca*. Altiarchaeum with its episymbiont *Ca*. Huberiarchaeum present in CG^12–14^ compared to MSI, where the symbiont is absent^15^ (Fig. 1, Fig. S2) and *Ca*. Altiarchaea is heavily targeted by viruses^16^. We hypothesize that the differences in the expression profile are not only influenced by the differing chemical composition of the two ecosystems, but also by the presence of the episymbiont in Crystal Geyser. The two sampling sites, CG and MSI, are located in Utah, USA (N 38 56’ 18.125’’; W 110 8’ 7.389’’) and in Regensburg, Germany (N 48 59’ 8.999’’; O 12 7’ 38.459’’), respectively, and also differ in chemical composition of bio-processable ions and other molecules in their groundwater^12,17^ (summarized in Fig. 1). Particularly, the chemical composition of CG varies throughout the phases of eruption^12^. When comparing the ion composition of CG’s minor eruption phase that sources groundwater from the deepest intersected aquifer that is dominated by *Ca*. Altiarchaea^12,17^ to MSI, the latter sampling site appears to be rather limited in ion and nutrient availability (Fig. 1). MSI as a sulfidic spring with high levels of sulfide, has a three-fold lower sulfate concentration than CG. Interestingly, nitrate, sodium and potassium have also a lower abundance in MSI compared to CG, which might influence the necessity to form biofilms for nutrient retention and filtration from groundwater. This is in agreement with the finding that *Ca*. Altiarchaea are predominantly present as single cells in CG^12^ (Fig. 1, Fig. S1) and almost solely found as biofilms in MSI (Fig. 1, Fig. S2), where the cells interconnect with *hami*^15,16,18^. Irrespective of these physiological differences and site-specific horizontal diversification of *Ca*. Altiarchaea, their core genome is highly conserved across many deep subsurface sites^11^ rendering it the ideal genus for studying differential gene expression with respect to different environmental conditions.

**Figure 1.**
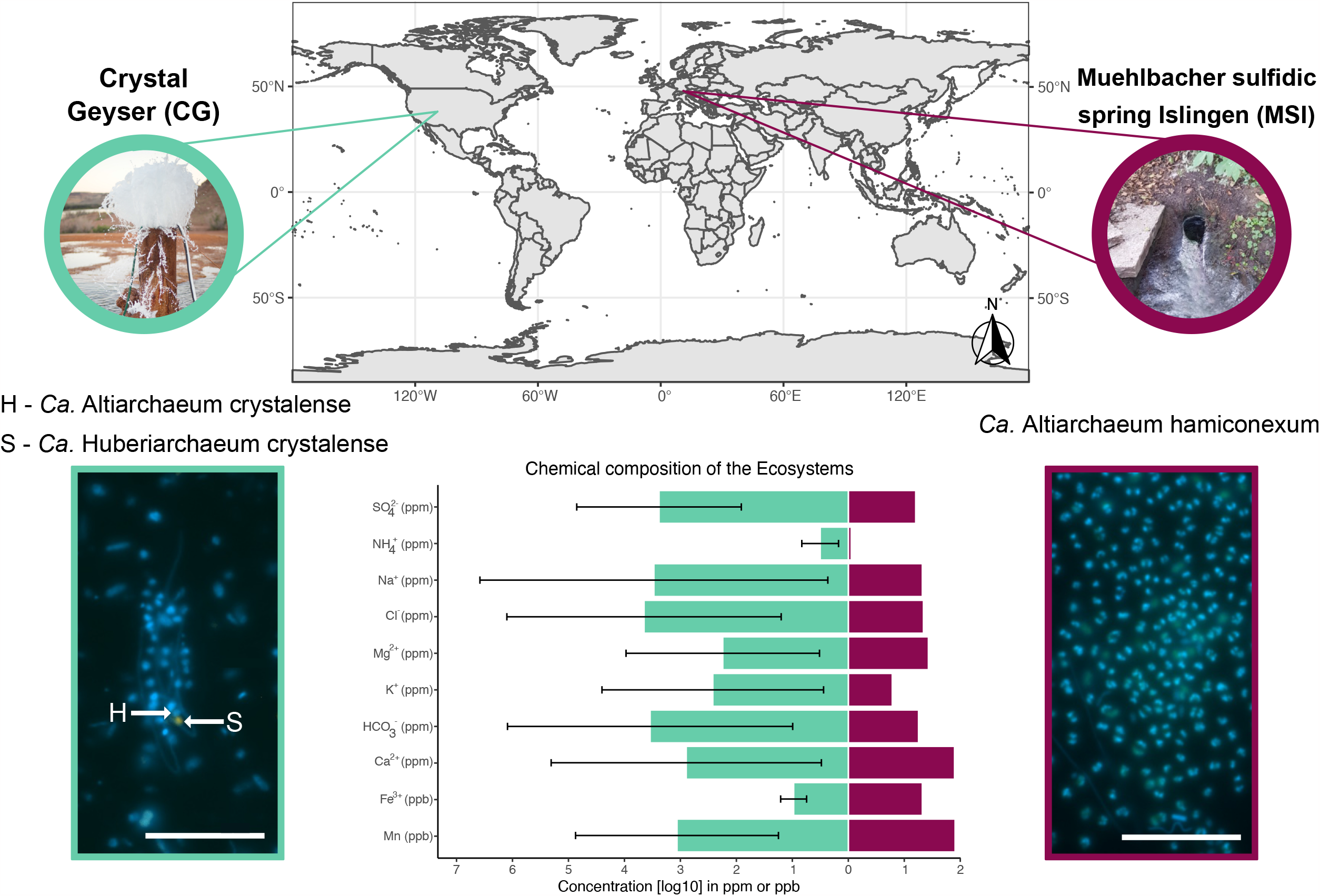
World map showing the two sampling sites, Crystal Geyser (CG, aquamarine) and Muehlbacher sulfidic spring (MSI, magenta). Differences in the chemical composition of MSI (n=1)^21^, and CG (n=3)^14^ are shown for all nutrients and ions measured in both ecosystems. Fluorescence *in-situ* hybridization (FISH) images show the host *Ca*. Altiarchaeum (SMARCH714 labeled with Atto448^17^) in green (H) and the symbiont *Ca*. Huberiarchaeum (HUB1206 with Cy3^13^) in orange (S). Shown chemical compositions from CG were sampled in the minor eruption phase of the eruption cycle, where *Ca*. Altiarchaea is the dominant organism. Please note that the symbiont was not detected in MSI. Scale bar 10 µm.

Predicted genes of previously published genomes from CG and MSI (Table S2) were clustered at 80% nucleotide similarity. Of the 537 shared gene clusters (Fig. 2A), 430 were assigned a functional annotation. The 107 remaining gene clusters were either not annotated (no hit in FunTaxDB 1.2, which is based on UniRef100 release 2023_02^19,20^) or annotated as uncharacterized proteins. From the 430 annotated gene clusters, 94 were within the first or last 200-bps of the respective scaffold causing irregularities in transcriptome mapping (underestimation of coverage). These genes were also excluded from downstream statistical analysis. The remaining 336 gene clusters were sorted according to their functional annotation, whereby the overall differential expression for most gene clusters in CG (∼90.8%, n=305) was higher than in MSI (∼9.2%, n=31).

**Figure 2.**
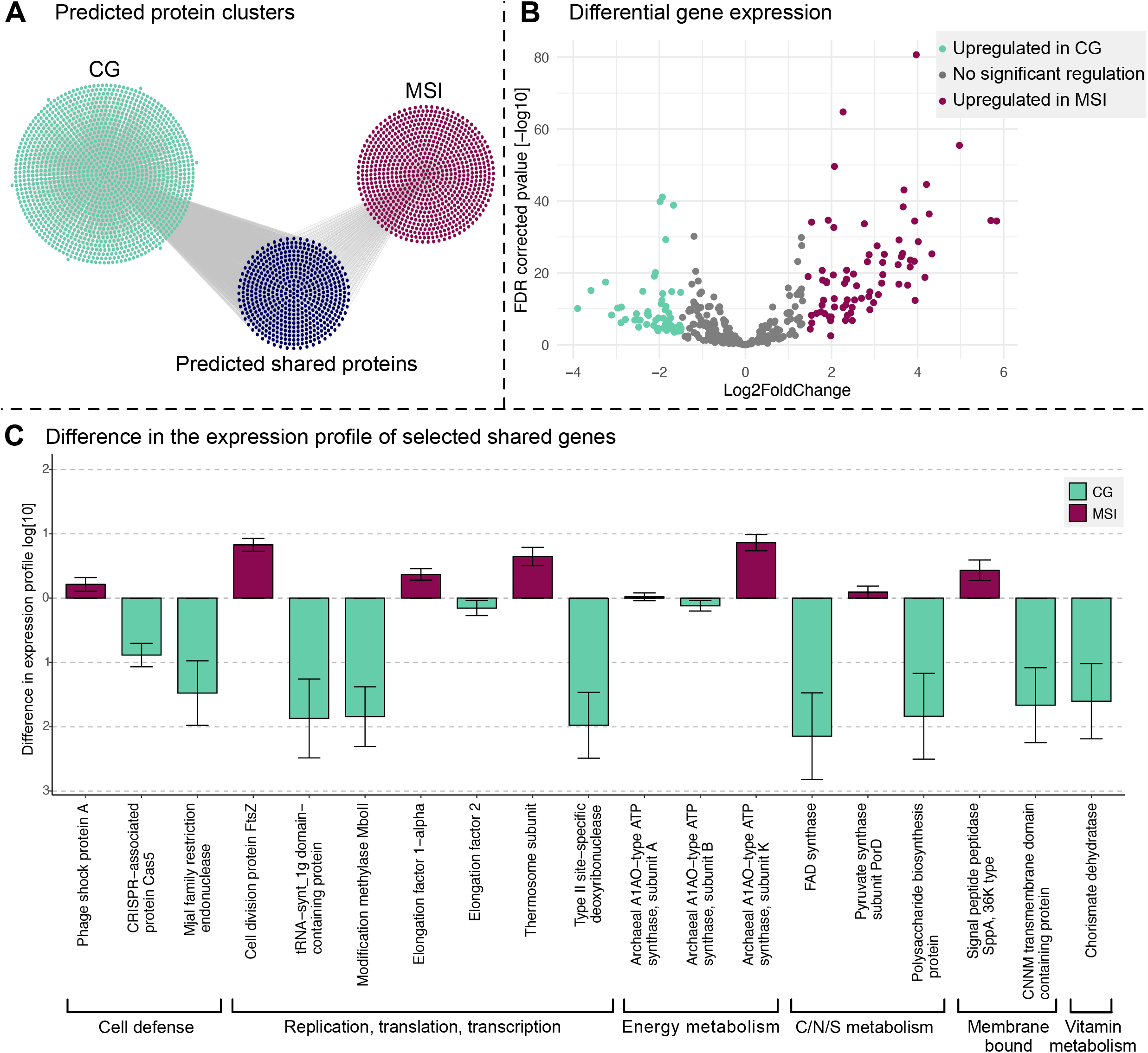
Gene clusters, differential gene expression and differences in gene expression profile of two *Ca*. Altiarchaea populations. **A** Gene clusters (80% AA similarity) of CG (n = 1447) and MSI (n = 824) as well as shared protein clusters (n = 537). **B** Differential gene expression of MSI compared to CG. The count data was normalized based on the coverage of ten ribosomal proteins (see methods) and then evaluated with DESeq2^31,32^ in R studio^33^. **C** Difference graph of the expression profile for 19 shared gene clusters selected based on annotation, difference, and relationship to physiology/ecology (the values represent the difference in the expression of the gene clusters). All values, including mean expression rate, and standard deviations are listed in Table S3.

Based on DESeq2 calculated Log2FoldChange (below -1.47 and above 1.44; FDR corrected p-values < 0.05), the overall regulation of genes revealed a significant upregulation of 76 genes in MSI compared to 54 genes in CG. Genes overexpressed in the latter ecosystem code for proteins that included Modification Methylase MboII, a Type II site-specific deoxyribonuclease, a FAD synthase, and an RNA synthase (Fig 2B). By contrast, MSI upregulated genes included those for ribosomal proteins (*e*.*g*., S11, S28e, L35Ae), a phage shock protein, a cell division protein (FtsZ), and a Thermosome subunit (Fig. 2B).

Focusing on the difference in the expression profile of shared gene clusters (Fig. 2C, Fig. S3-S10) it was evident that the increase in expression of any given significantly differently expressed gene was substantially greater in CG than in MSI (Fig. S11-S12). Particularly, some genes related to replication (*e*.*g*., DNA polymerases, Table S3-S4) and nutrient metabolism have an up to two-fold higher expression in CG than in MSI (Fig S5 and S8) suggesting that *Ca*.

A. crystalense is more active and replicating than *Ca*. A. hamiconexum when sampling the respective ecosystem. However, the expression of FtsZ, a protein involved in forming the septum of a dividing *Ca*. A. hamiconexum^15,16,18^ cell seems upregulated in MSI (Fig. 2C). The accumulation of this protein, which usually has similar concentrations in the cell irrespective of cell division, can be used as an indicator of a starting cell division process as it localizes as mid-cell ring early in the division process shown for *E. coli*^21^. In addition, genes for the elongation factor 1 alpha, which is included in the aminoacyl tRNA incorporation in archaea and eukaryotes^22^, and the thermosome subunit, which represents the chaperonin family in archaea and is accordingly involved in the protein folding^23,24^, are up to two fold higher expressed in MSI as compared to CG. This can also be an indicator that the cell division in MSI is upregulated. Therefore, the upregulation of the ftsZ gene suggests a higher replication rate of *Ca*. Altiarchaeum hamiconexum in MSI and supports the visible diploidy of the cells in previously published^15,25^ and here shown fluorescence *in situ* hybridization (FISH) images (Fig. 1; Fig S1-S2). Comparing the function of the cellular replication genes upregulated in CG (*i*.*e*., genes encoding DNA polymerases) versus MSI (*i*.*e*., genes encoding for the cell division process), it is indicated that *Ca*. A. crystalense appears to heavily replicate the genome but somehow does not proceed the cell division cycle as corresponding cells in MSI. Previous studies on archaeal cell division showed that the depletion of the FtsZ proteins in *Haloferax volcanii* inhibits cell division, while DNA replication is still ongoing^26^. We propose that although the DNA synthesis is very prominent in Altiarchaea from CG, their final cell division seems hampered, likely due to the presence of the symbiont *Ca*. H. crystalense or many different viruses in CG that show infection histories with this organism^14^.

*Ca*. A. hamiconexum has been described to be infected by at least two different viruses, one of which is a lytic virus^16,27^. In agreement with these findings, the differential expression analysis revealed a significant increase of the gene encoding for a phage shock proteins, which are a stress response to membrane penetration by invading mobile genetic elements (MGEs)^28^, in MSI compared to CG. By contrast, CRISPR Cas 5, which is a protein involved in the cascade building for the splicing mechanisms in CRISPR type I systems (reviewed by Hille and Carpentier, 2016)^29^ is upregulated in *Ca*. A. crystalense (Fig. S2C), which shows an expansive spacer variety over six years not only against MGEs but also against its episymbiont^30^. Consequently, we identified a specific adaptation of defense mechanisms against lytic viruses and the episymbiont, respectively.

Beyond upregulation of the CRISPR system, the interaction of the episymbiont might also be responsible for the upregulation of other metabolic functions in *Ca*. A. crystalense. For example, multiple genes encoding for proteins involved in energy metabolism such as the quinolinate synthase, FO synthase subunit 2 and the FAD synthase (Fig. 2C, Fig. S5-S6) were significantly enriched in the CG transcriptome. Increased energy demands might stem from the highly active CRISPR Cas system which acquired hundreds of thousands of different spacers in the *Ca*. Altiarchaea population^14^. In addition, polysaccharide biosynthesis genes were found to be upregulated in CG, although we only seldomly found biofilms of *Ca*. Altiarchaea in CG compared to MSI (Fig. 1). This upregulation could either be related to the scavenging nature of the episymbiont^13^ or indicate an intrinsic tendency of the *Ca*. A. crystalense to form biofilms without success potentially due to the turbulent geyser system^12^. In summary, we found that gene expression of two *Ca*. Altiarchaea populations, originating from distinct geological settings, is influenced by environmental factors and biological interactions. While the upregulated genes in the microbial population in MSI appear reflective of viral attacks, the episymbiont in CG and the turbulence of the geyser system seem to upregulate the CRISPR system, the energy metabolism, and the biofilm formation. Leveraging metatranscriptomes of low-biomass deep subsurface ecosystems, this study contributes to the existing DNA sequencing-based body of literature on the deep biosphere in general and on *Ca*. Altiarchaea in particular.

## Supporting information

Esser_et_al_Supplementary_Information

## ACKNOWLEDGEMENTS

This effort was funded by the Ministerium für Kultur und Wissenschaft des Landes Nordrhein-Westfalen (“Nachwuchsgruppe Dr. Alexander Probst”) and the German Research Foundation under project NOVAC (grant number DFG PR1603/2-1). The work (proposals: 10.46936/10.25585/60000685, 10.46936/10.25585/60000800) conducted by the U.S.

Department of Energy Joint Genome Institute (https://ror.org/04xm1d337), a DOE Office of Science User Facility, is supported by the Office of Science of the U.S. Department of Energy operated under Contract No. DE-AC02-05CH11231. JP was supported by Aker BP within the framework of the GeneOil Project given to AJP. The sequencing experiments presented in this paper were carried out at the LCSB sequencing platform (RRID: SCR_021931) at the University of Luxembourg. We thank Christopher T Brown (Metagenomi) for providing the image of Crystal Geyser, Ken Dreger for exemplary server administration, and Sabrina Eisfeld, Ines Pothmann, and Maximilliane Ackers for administrative support.

## CONFLICT OF INTEREST

The authors declare no conflict of interest.

### Data availability statement

All metatranscriptomics datasets are published under the accession number SAMN14515498, SAMN14515403, SAMN14515402 for CG and under the BioProject number PRJNA1005487 with the individual Accession numbers SRX21390066, SRX21390067, and SRX21390068 for MSI. The accession numbers of metagenomic assembled genomes are listed in Table S1.

### Author Contribution

SPE performed RNA extraction for MSI and bioinformatical analysis of this study. VT did FISH imaging with assistance of IB. JL performed RNA extractions for CG. JP, CM and AJP helped with bioinformatical assistance. SPE and AJP conceptualized the study. The manuscript was written by SPE and AJP with input from all co-authors.

