## Supplementary material for "Differential expression of core metabolic functions in *Candidatus* Altiarchaeum inhabiting distinct subsurface ecosystems": Esser_et_al_Supplementary_Information

Content:

1. Methods
2. Supplementary Figures
3. Supplementary Tables
4. List of additional supplementary Data
5. References

### 1. Methods

#### RNA extraction and sequencing

For RNA extraction from *Ca.* Altiarchaeota biofilms in MSI the biofilm flocks were harvested in November 2021 as previously described[1] and directly frozen at - 80°C until further processing in the lab. RNA was extracted with the RNeasy PowerBiofilm RNA extraction kit (Qiagen, Germany) according to the manufacturer's instructions. The extracted RNA was sequenced at the LCSB (Luxemburg) with 150bp paired-end Illumina technology. Prior to sequencing, the rRNA was depleted using Zymo-Seq RiboFree Total RNA Library Kit (R3003) according to the manufacturer protocol. The library amplification was performed with 12 PCR cycles.

Sequencing data from Crystal Geyser was retrieved from a previous study[2] (minor eruption phase, samples CG05, CG08, and CG16), specifically using datasets from samples where *Ca.* Altiarchaeum crystalense is the most abundant organism. In brief, the erupted water from CG was sequentially filtered on PTFE filters, which were frozen on site on dry ice[3]. The RNA extraction was performed with an adjusted protocol of the Qiagen DNeasy PowerMax Soil kit (Qiagen, Germantown, MD)[2]. All accession numbers are listed in Table S1.

#### Coverage-based normalization of metatranscriptomes

After quality filtering with bbduk (https://github.com/BioInfoTools/BBMap/blob/master/sh/bbduk.sh) and sickle[4], the metatranscriptomics reads were normalized by mapping[5] reads against representative genomes of *Ca.* Altiarchaeum hamiconexum (MSI) and *Ca.* A. crystalense (CG) (see Table S1 for accession numbers). The mean coverage of ten house-keeping genes [30S ribosomal proteins: S4, S5, S7, S8e, S9, S10, S11, S12, S13, S15] were used to calculate the normalization factor of each metatranscriptome sample (per ecosystem n=3) [Table S2]. Prior to choosing these genes for normalization the position of the gene on the scaffold (not within the first/last 200 bp of the scaffold) and a stable coverage distribution across the gene was taken into consideration.

#### Clustering of genes and calculation of expression profile

Genes of the abovementioned *Ca.* A. hamiconexum and *Ca.* A. crystalense genomes were predicted with prodigal[6], and consecutively clustered with cdhit[7,8] at 80% amino acid similarity. The genes from the two ecosystems sharing a cluster were annotated with the FunTaxDB[9] (version 1.2) database. All genes that either had no or an unclassified annotation and/or which started/ended within the first/last 200 bps of the scaffolds were discarded within the expression profile. Removing genes starting/ending within the first/last 200 bps of a scaffold was chosen to avoid mapping distortions resulting from inaccurate mapping at these scaffold regions.

The abovementioned normalization factors were used to determine the coverage differences introduced by the varying extraction and sequencing methods and the mean coverage with standard deviation of the shared gene clusters were calculated. The visualization was performed with ggplot2[10] in R studio[11,12] (version 2023.03.0+386). The data for evaluating the count data within the RNA-seq data was performed with DESeq2 implemented in R[13]. The threshold values of up- and downregulation of differential gene expression were calculated to the base expression of MSI according to Quackenbush 2002[14].

**Fluorescence *in-situ* hybridization (FISH)**

*Ca.* A. hamiconexum biofilm flocks were taken from MSI. For FISH purposes, the biofilm flocks were harvested and fixed as previously described on site with 3% (v/v)% formaldehyde[15], stored at -20 °C, and deposited on a slide for hybridization. For *Ca.* A. crystalense (CG), cells were filtered onto 0.2 µm PTFE filters (Polytetrafluorethylene), and fixed on site with 3% (v/v)% formaldehyde and stored at -80°C as described previously[16]. The samples from CG and MSI were taken in August 2021 and February 2022, respectively. For both systems the 16S rRNA probe “SMARCH714” (Moissl et al. 2003)[17] was labeled with Atto488. The episymbiont *Ca.* Huberiarchaeum crystalense was labeled with a specifically designed Cy3 probe called “HUB1206”[16]. The FISH procedure was carried out as previously described by Schwank et al.[16]. Cells were counterstained with DAPI (4 µg $mL^{-1}$).

Image analysis was carried out with a Zeiss Axio Imager M2m epifluorescence microscope (X-Cite XYLIS Broad Spectrum LED Illumination System, Excelitas) equipped with an Axio Cam MRm and a Zen 3.4 Pro software (version 3.4.91.00000). Imaging was carried out by using the 100x/1.3 oil objective EC-Plan NEOFLUAR and three different filter sets: 09 for achieving the 16S rRNA signals of *Ca.* Altiarchaea, 43 Cy3 for the detection of *Ca.*

Huberiarchaea signals, and 49 DAPI for imaging *Ca.* A. crystalense/hamiconexum cells and *Ca.* H. crystalense cells.

### 2. Supplementary Figures

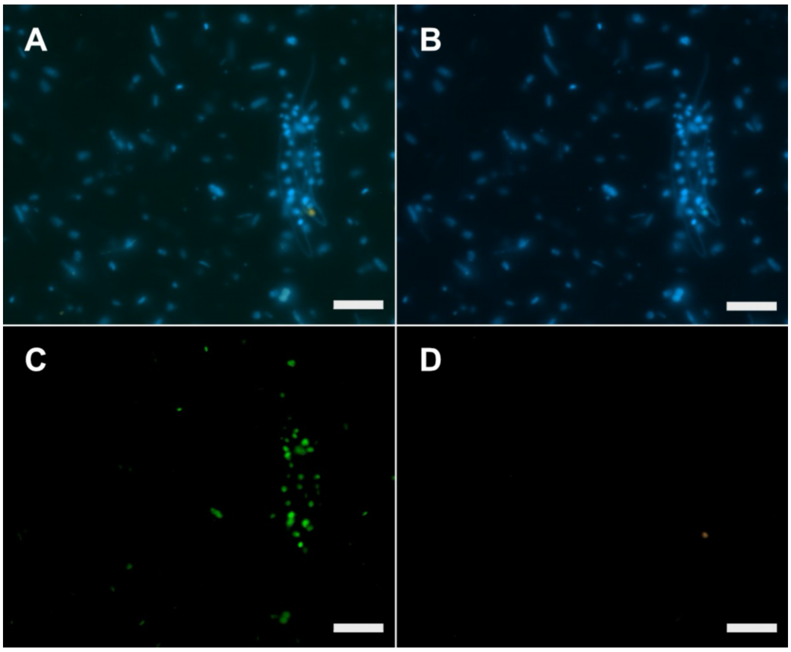


**Figure S1|** Fluorescence *in situ* hybridization images of *Ca.* Altiarchaeum crystalense and *Ca.* Huberiarchaeum crystalense in Crystal Geyser. **A** Merged FISH images as shown in Fig. 1. **B** DAPI staining of the sample. **C** 16S rRNA of *Ca.* Altiarchaeum labeled using the SMARCH714 probe[17] with Atto488. **D 16S rRNA of** *Ca.* Huberiarchaeum crystalense labeled using the Hub1206 probe[16] and Cy3. Scale bar = 10 µm.

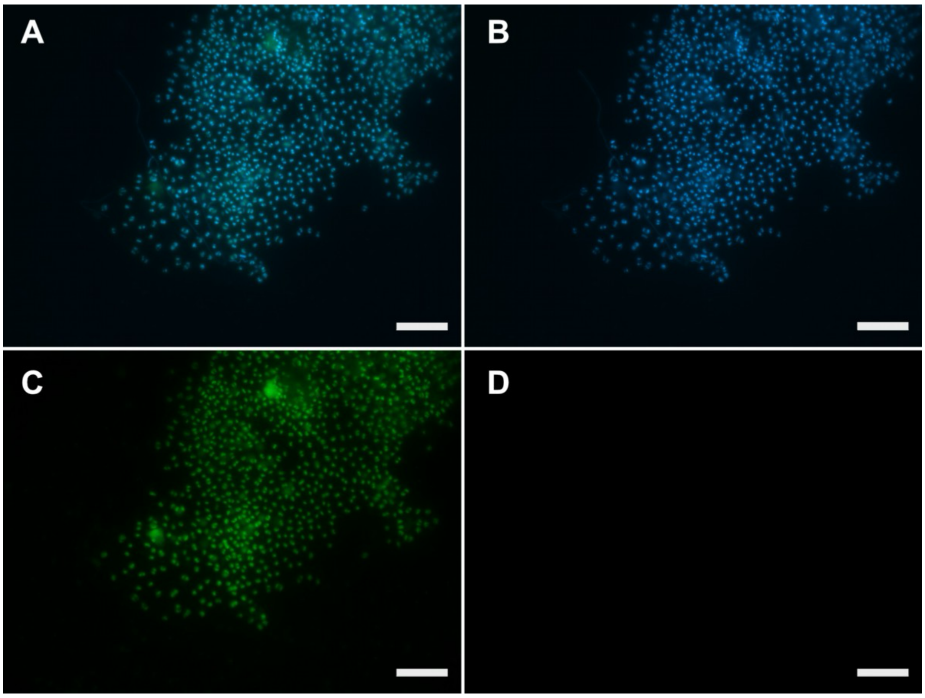


**Figure S2|** Fluorescence *in situ* hybridization images of *Ca.* Altiarchaeum hamiconexum **A** Merged FISH images as shown in Fig. 1. **B** DAPI staining of the sample. **C** 16S rRNA of *Ca.* Altiarchaeum labeled using the SMARCH714 probe[17] with Atto488. **D 16S rRNA of** *Ca.* Huberiarchaeum crystalense labeled using the Hub1206 probe[16] and Cy3. Scale bar = 10 µm.

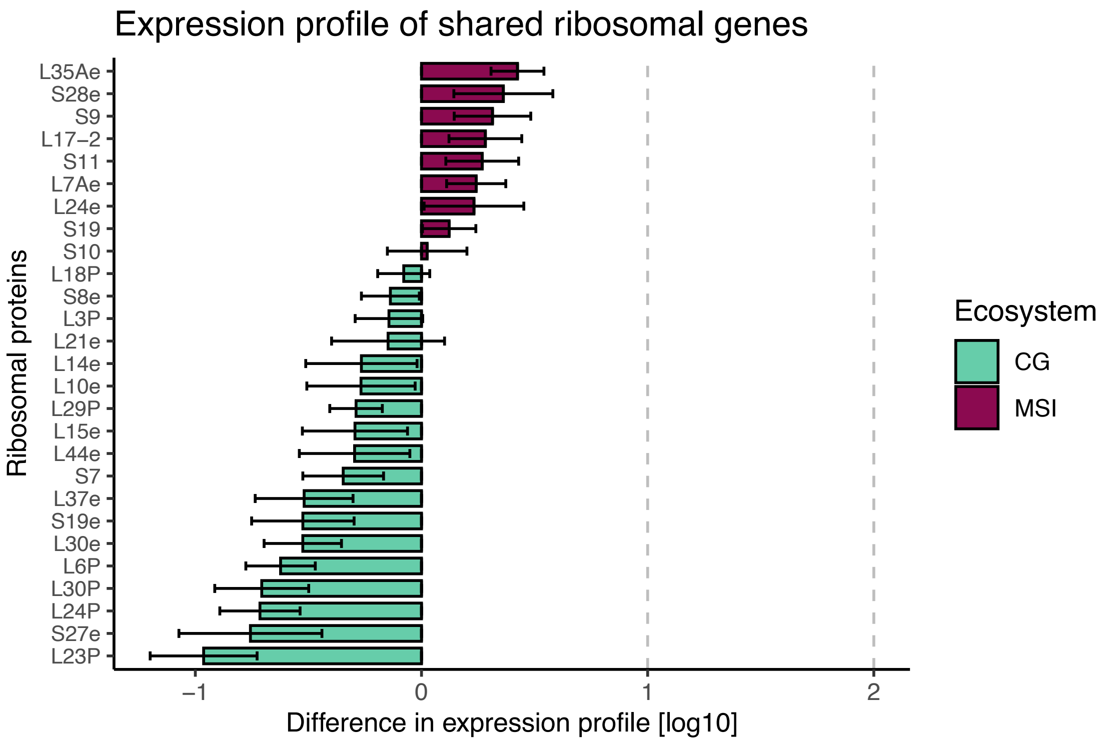


**Figure S3|** Difference in expression profile of ribosomal proteins derived from metatranscriptomics data of CG (blue) and MSI (magenta), respectively. Visualization was performed with ggplot2[10] in R studio[11,12].

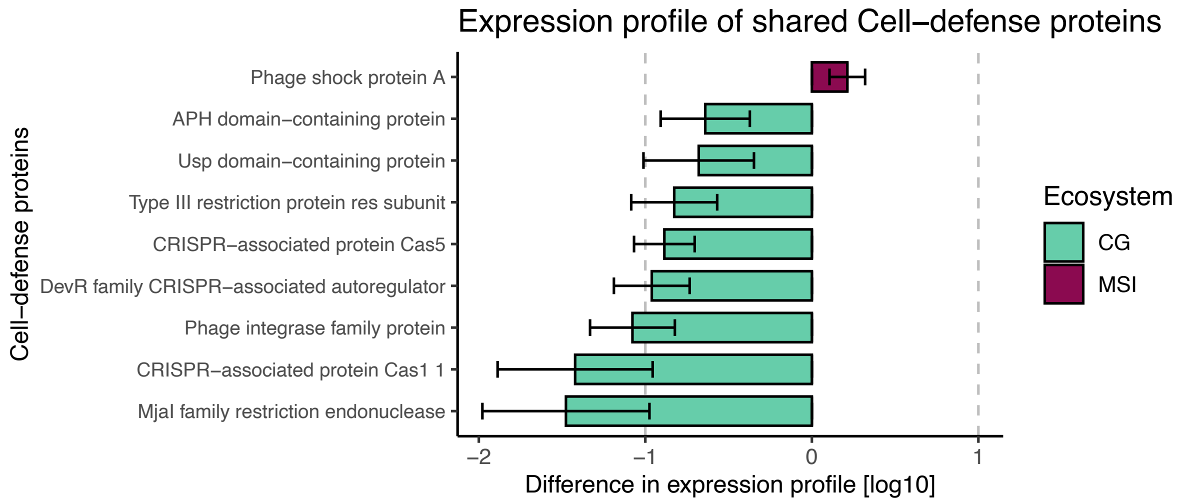


**Figure S4|** Difference in expression profile of proteins related to cell-defense and derived from metatranscriptomics data of CG (blue) and MSI (magenta), respectively. Visualization was performed with ggplot[10] in R studio[11,12].

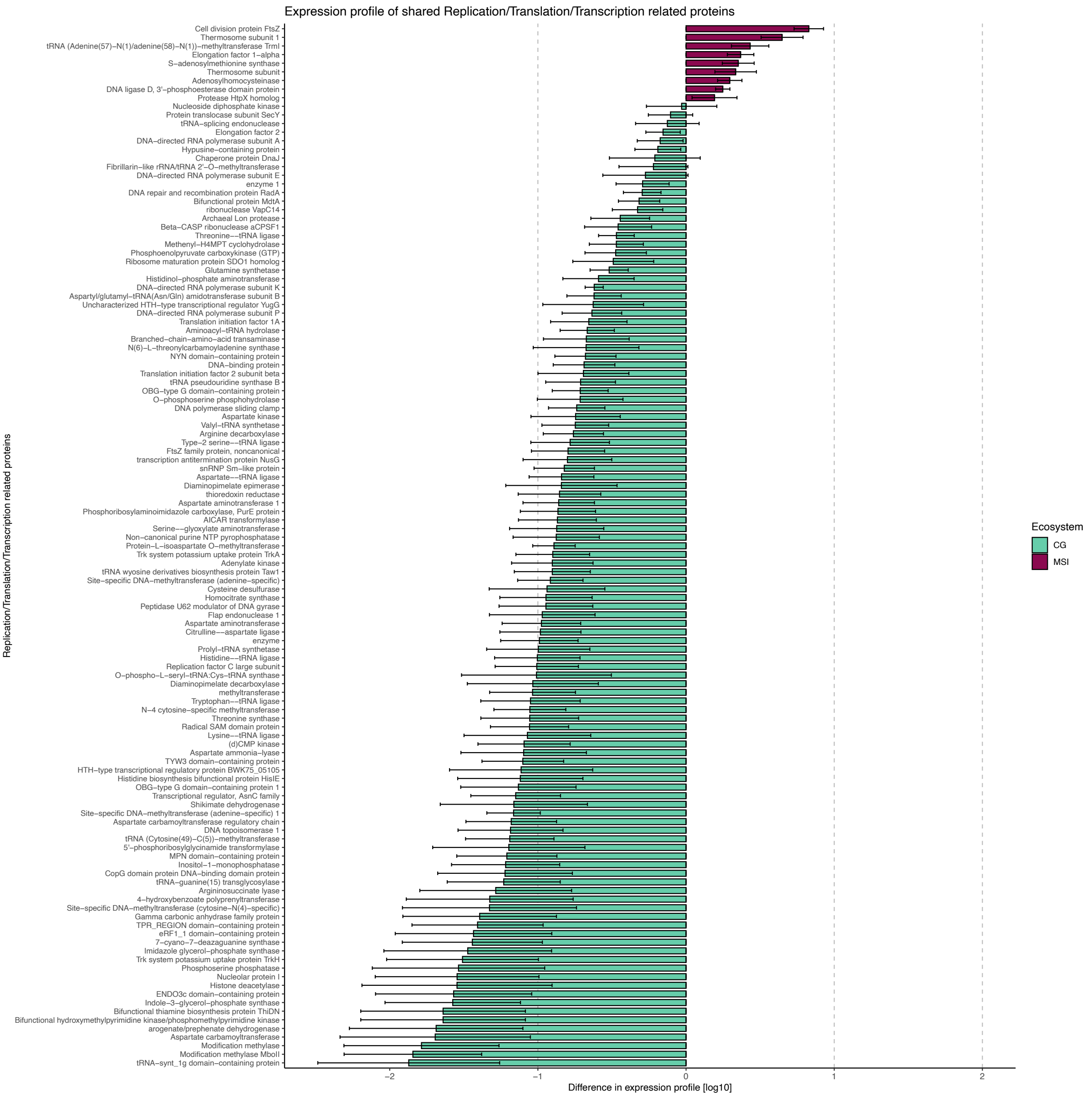


**Figure S5|** Difference in expression profile of replication, translation and transcription related proteins derived from metatranscriptomics data of CG (blue) and MSI (magenta). Visualization was performed with ggplot[10] in R studio[11,12].

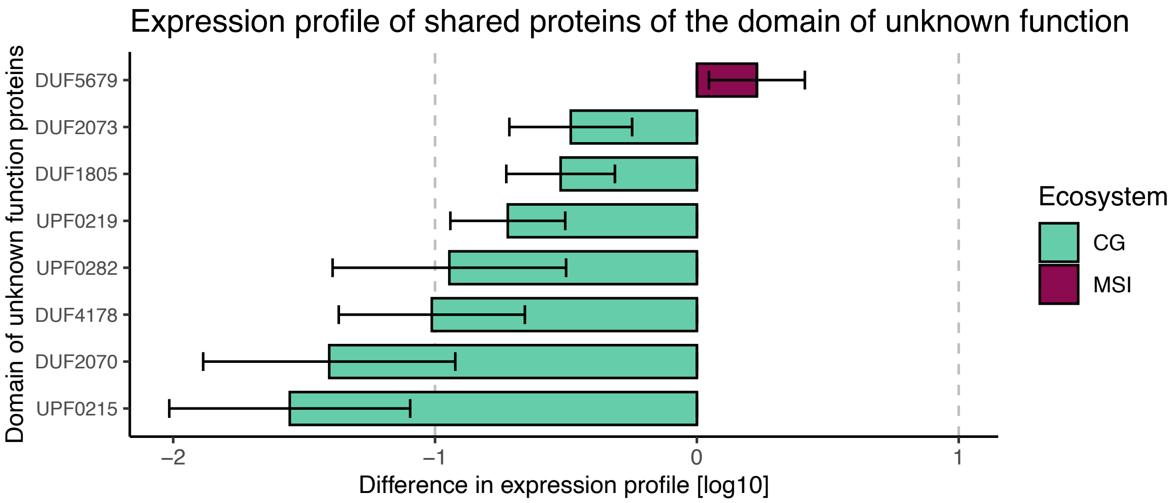


**Figure S6|** Difference in expression profile of proteins classified as "domain of unknown function" and derived from metatranscriptomics data of CG (blue) and MSI (magenta). Visualization was performed with ggplot[10] in R studio[11,12].

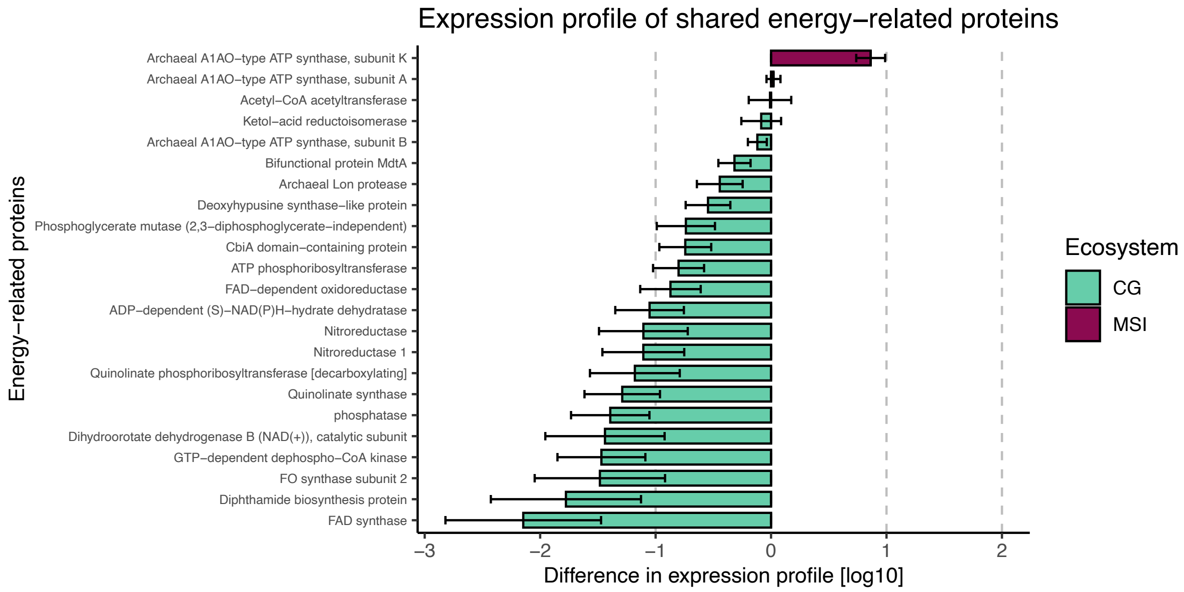


**Figure S7|** Difference in expression profile of energy related proteins derived from metatranscriptomics data of CG (blue) and MSI (magenta). Visualization was performed with ggplot[10] in R studio[11,12].

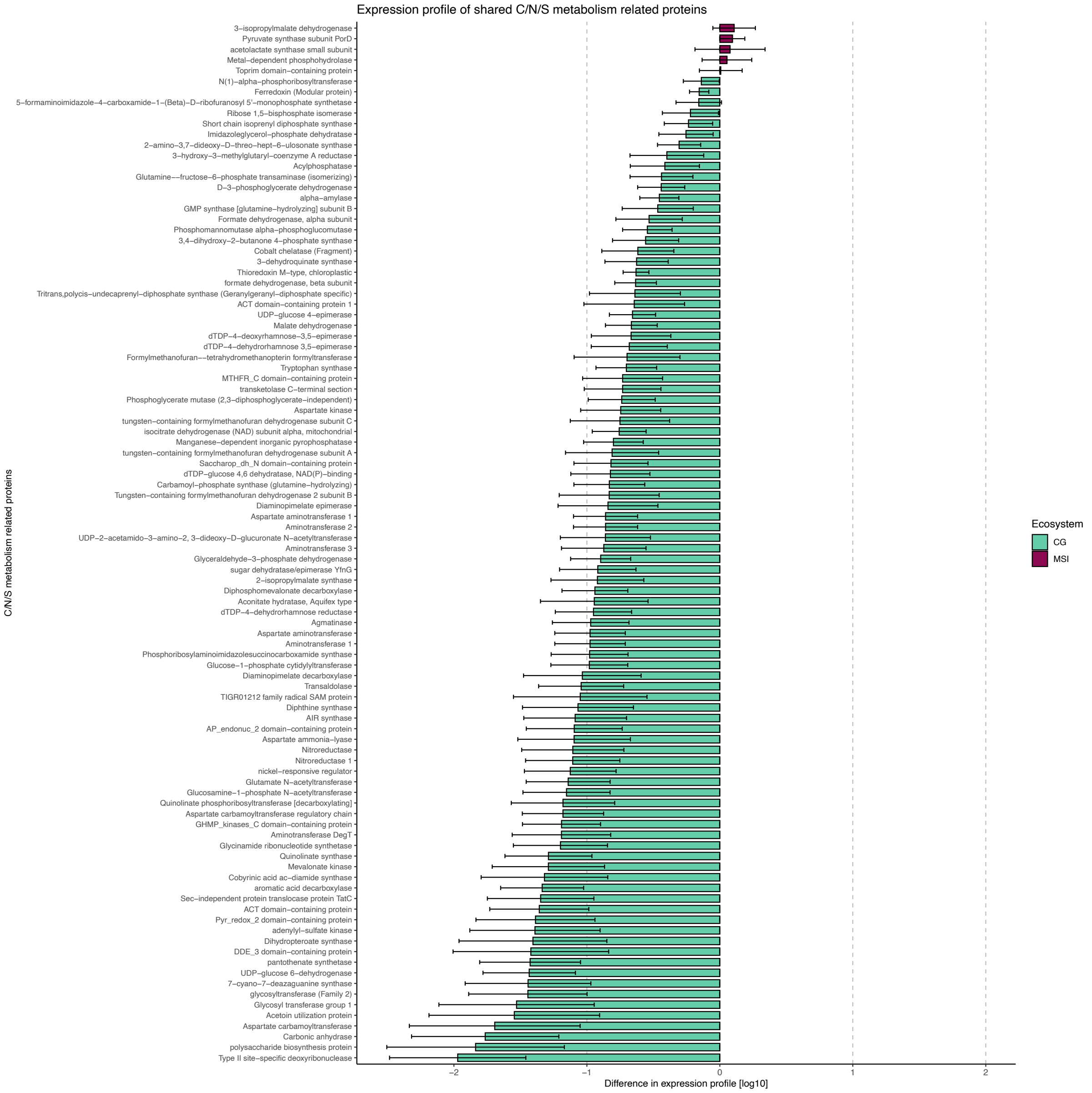


**Figure S8|** Difference in expression profile of carbon, nitrogen and sulfur metabolism related proteins derived from metatranscriptomics data of CG (blue) and MSI (magenta). Visualization was performed with ggplot[10] in R studio[11,12].

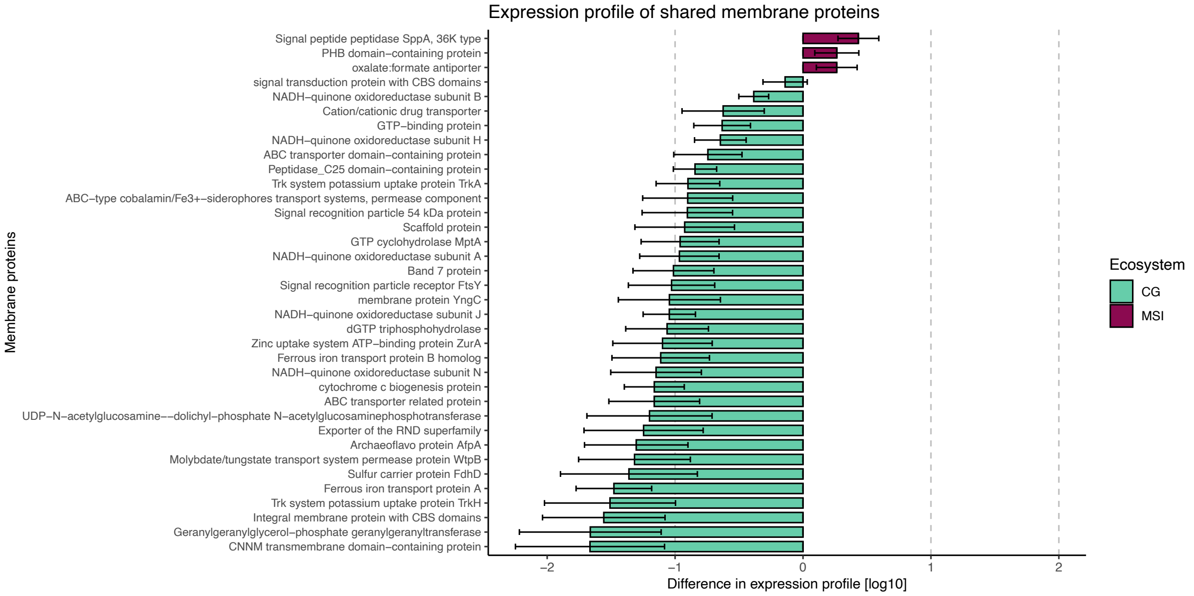


**Figure S9|** Difference in expression profile of membrane proteins derived from metatranscriptomics data of CG (blue) and MSI (magenta). Visualization was performed with ggplot[10] in R studio[11,12].

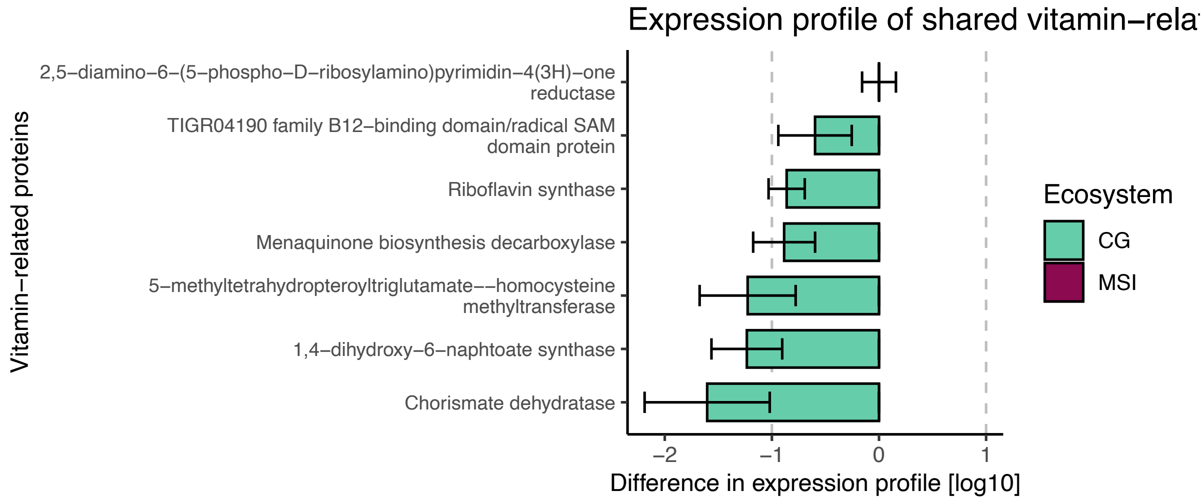


**Figure S10|** Difference in expression profile of vitamin related proteins derived from metatranscriptomics data of CG (blue) and MSI (magenta). Visualization was performed with ggplot[10] in R studio[11,12].

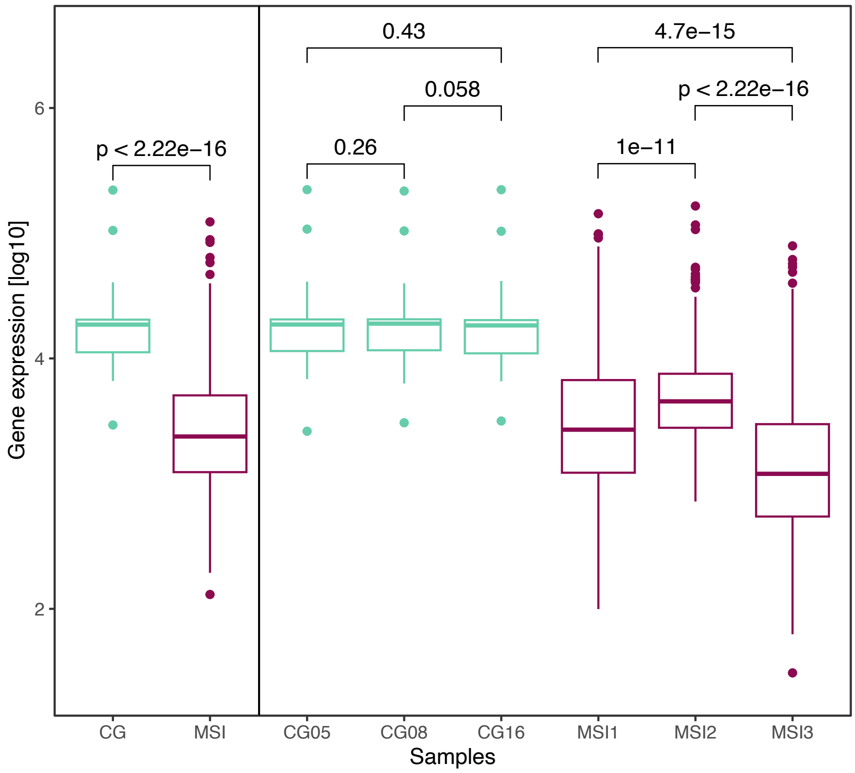


**Figure S11|** Inter and intra ecosystem comparison of the mean normalized gene expression of CG's and MSI's shared gene clusters (Kruskal-Wallis Anova, *p*-values within figure). CG showed a significantly higher expression of the core metabolic pathways then MSI. Data visualization was performed with ggplot[10] in R studio[11,12].

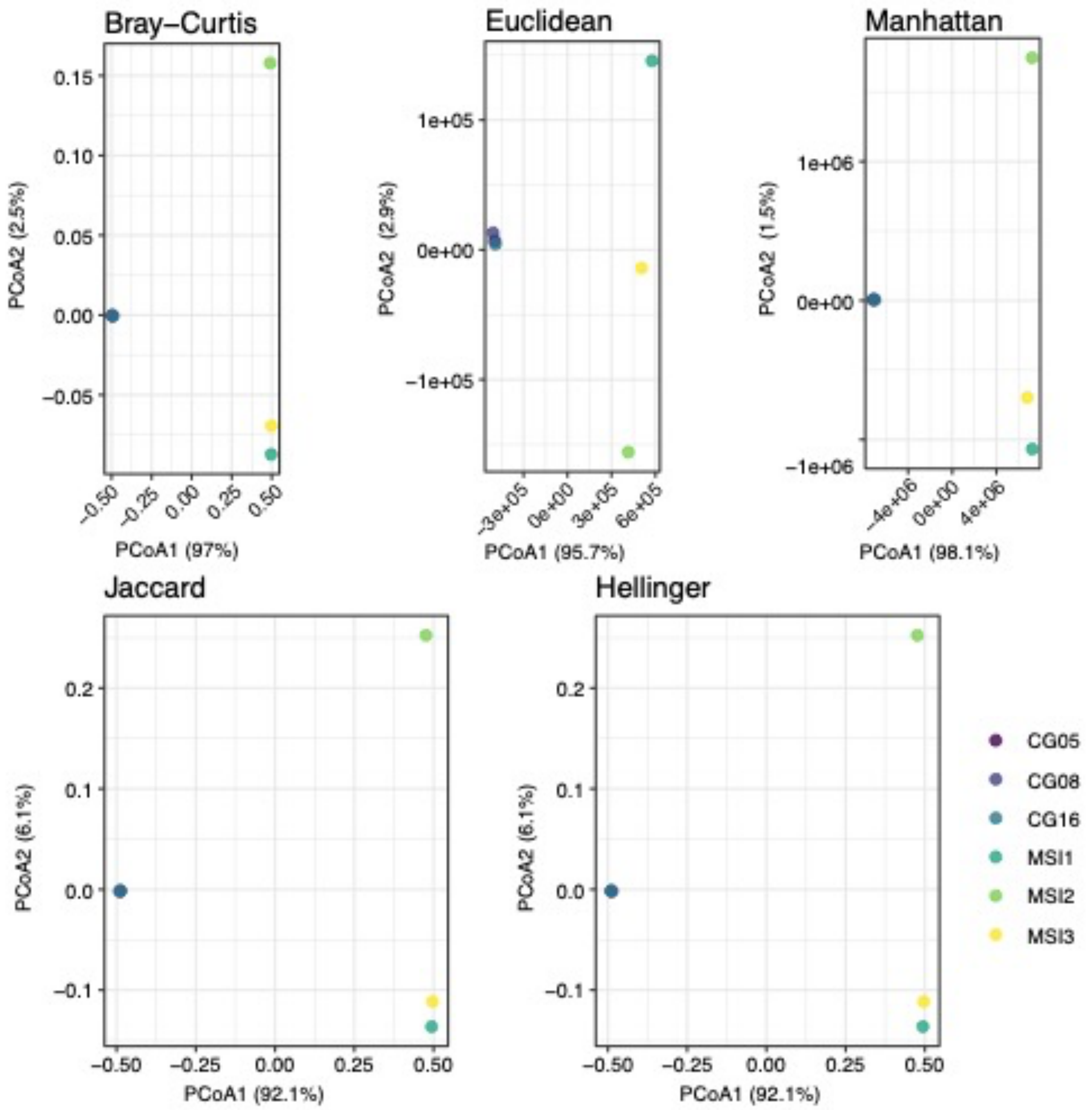


**Figure S12|** Principal Coordinate Analyses (PCoA) of normalized gene expression within the metatranscriptomic dataset. Compared are different calculations of the dissimilarity matrix. All variants of the PCoAs show that the datasets of Crystal Geyser and Muehlbacher sulfidic spring cluster separated from each other. Data visualization was performed with ggplot[10] in R studio[11,12].

### 3. Supplementary Tables

**Table S1|** Accession Numbers and metadata of metatranscriptomics reads and *Ca.* Altiarchaea genomes used in this study

| | Type of data | Ecosystem | Abbr./Genome name | Accession number | Year |
|---|---|---|---|---|---|
| **1** | Reads | CG | CG05 | SAMN14515498 [NCBI] | 2015 |
| **2** | Reads | CG | CG08 | SAMN14515403 [NCBI] | 2015 |
| **3** | Reads | CG | CG16 | SAMN14515402 [NCBI] | 2015 |
| **4** | Reads | MSI | MSI1 | SRX21390066 [NCBI] | 2021 |
| **5** | Reads | MSI | MSI2 | SRX21390067 [NCBI] | 2021 |
| **6** | Reads | MSI | MSI3 | SRX21390068 [NCBI] | 2021 |
| **7** | Genome | CG | Candidatus Altiarchaeum sp. CG_4_8_14_3_um_filter_33_2054 | 2786546689 [JGI IMG] | 2014 |
| **8** | Genome | CG | Candidatus Altiarchaeum sp. CG_4_9_14_0_8_um_filter_32_206 | 2786546690 [JGI IMG] | 2014 |
| **9** | Genome | CG | Candidatus Altiarchaeum sp. CG_4_10_14_0_8_um_filter_32_851 | 2786546691 [[JGI IMG] | 2014 |
| **10** | Genome | CG | Candidatus Altiarchaeum sp. CG2_30_32_3053 | 2786546688 [JGI IMG] | 2009 |
| **11** | Genome | CG | Candidatus Altiarchaeum sp. CG03_land_8_20_14_0_80_32_618 | 2786546693 [JGI IMG] | 2014 |
| **12** | Genome | CG | Candidatus Altiarchaeum sp. CG12_big_fil_rev_8_21_14_0_65_33_22 | 2786546692 [JGI IMG] | 2014 |
| **13** | Genome | CG | Candidatus Altiarchaeum sp. CG1_02_FULL | https://doi.org/10.6084/m9.figshare.22339555.v1 | 2009 |
| **14** | Genome | CG | Candidatus Altiarchaeum sp. CG1_02_FULL_1 | https://doi.org/10.6084/m9.figshare.22339555.v1 | 2009 |
| **15** | Genome | CG | Candidatus Altiarchaeum sp. CG2_30_FULL | https://doi.org/10.6084/m9.figshare.22339555.v1 | 2009 |
| **16** | Genome | CG | Candidatus Altiarchaeum sp. CG2_30_SUB10 | https://doi.org/10.6084/m9.figshare.22339555.v1 | 2009 |
| **17** | Genome | CG | Euryarchaeota archaeon JGI CrystG Aug3-3-F14 | 2693430074 [JGI IMG] | 2014 |
| **18** | Genome | CG | Euryarchaeota archaeon JGI CrystG Aug3-1-I17 | 2693430078 [JGI IMG] | 2014 |
| **19** | Genome | CG | Euryarchaeota archaeon JGI CrystG Aug3-3-C16 | 2693430067 [JGI IMG] | 2014 |
| **20** | Genome | CG | Euryarchaeota archaeon JGI CrystG Aug3-1-C22 | 2693430068 [JGI IMG] | 2014 |
| **21** | Genome | CG | Euryarchaeota archaeon JGI CrystG Aug3-3-I21 | 2693430072 [JGI IMG] | 2014 |
| **22** | Genome | CG | Euryarchaeota archaeon JGI CrystG Aug3-3-I7 | 2693430046 [JGI IMG] | 2014 |
| **23** | Genome | CG | Euryarchaeota archaeon JGI CrystG Aug3-1-F4 | 2693430045 [JGI IMG] | 2018 |
| **24** | Genome | MSI | Candidatus Altiarchaeum hamiconexum | SAMN18220766 [NCBI] | 2014 |

### 4. List of supplementary tables

**Table S2 |** Normalization factors calculated based on ten ribosomal proteins that are shared at 80% amino acid similarity in CG and MSI.

**Table S3 |** Raw and normalized coverages of shared gene clusters in CG and MSI. Normalization was done based in the in Table S2 calculated normalization factors.

**Table S4 |** Gene annotations of shared gene clusters in CG and MSI based the FunTaxDB 1.2. [accessed September 2022]
